# Antibody evasiveness of SARS-CoV-2 subvariants KP.3.1.1 and XEC

**DOI:** 10.1101/2024.11.17.624037

**Authors:** Qian Wang, Yicheng Guo, Ian A. Mellis, Madeline Wu, Hiroshi Mohri, Carmen Gherasim, Riccardo Valdez, Lawrence J. Purpura, Michael T. Yin, Aubree Gordon, David D. Ho

**Author notes:** These authors contributed equally (Qian Wang, Yicheng Guo, Ian A. Mellis, Madeline Wu). Correspondence (D.D.H.).

## Abstract

SARS-CoV-2 continues to evolve and spread around the world, and it remains critical to understand the functional consequences of mutations that lead to dominant viral variants. KP.3.1.1 is currently the most prevalent subvariant worldwide, while the recombinant subvariant XEC is exhibiting the fastest growth rate. Here we measured the in vitro neutralization of KP.3.1.1 and XEC by human sera, monoclonal antibodies, and soluble hACE2 receptor relative to their parental subvariants KP.3 and JN.1. KP.3.1.1 and XEC were slightly more resistant (1.3-1.6-fold) than KP.3 to serum neutralization, and the resultant antigenic map showed that the new subvariants are antigenically similar. Both also demonstrated greater resistance to neutralization by select monoclonal antibodies and soluble hACE2, all of which target the top of the viral spike. Our findings suggested that upward motion of the receptor-binding domain in spike is partially hindered by the N-terminal-domain mutations found KP.3.1.1 and XEC, thereby allowing these subvariants to better evade serum antibodies that target the viral spike when it is in the up position and thus having a growth advantage in the population.

## Main Text

The SARS-CoV-2 Omicron JN.1 subvariant rapidly increased in prevalence around the world since late 2023, and its progeny sublineages KP.2 and KP.3 were dominant successively and briefly thereafter. Recently, KP.3.1.1, bearing S31 deletion (S31Δ) on top of the KP.3 spike, has become the most prevalent subvariant worldwide^1^ (**Figure 1A**). XEC, a recombinant of JN.1 subvariants KS.1 and KP.3.3, is now showing a growth advantage in the population. XEC carries two additional spike N-terminal domain (NTD) mutations T22N and F59S beyond those in KP.3 (**Figure 1B**). How these spike mutations confer a growth advantage to KP.3.1.1 and XEC remains unknown.

**Figure 1.**
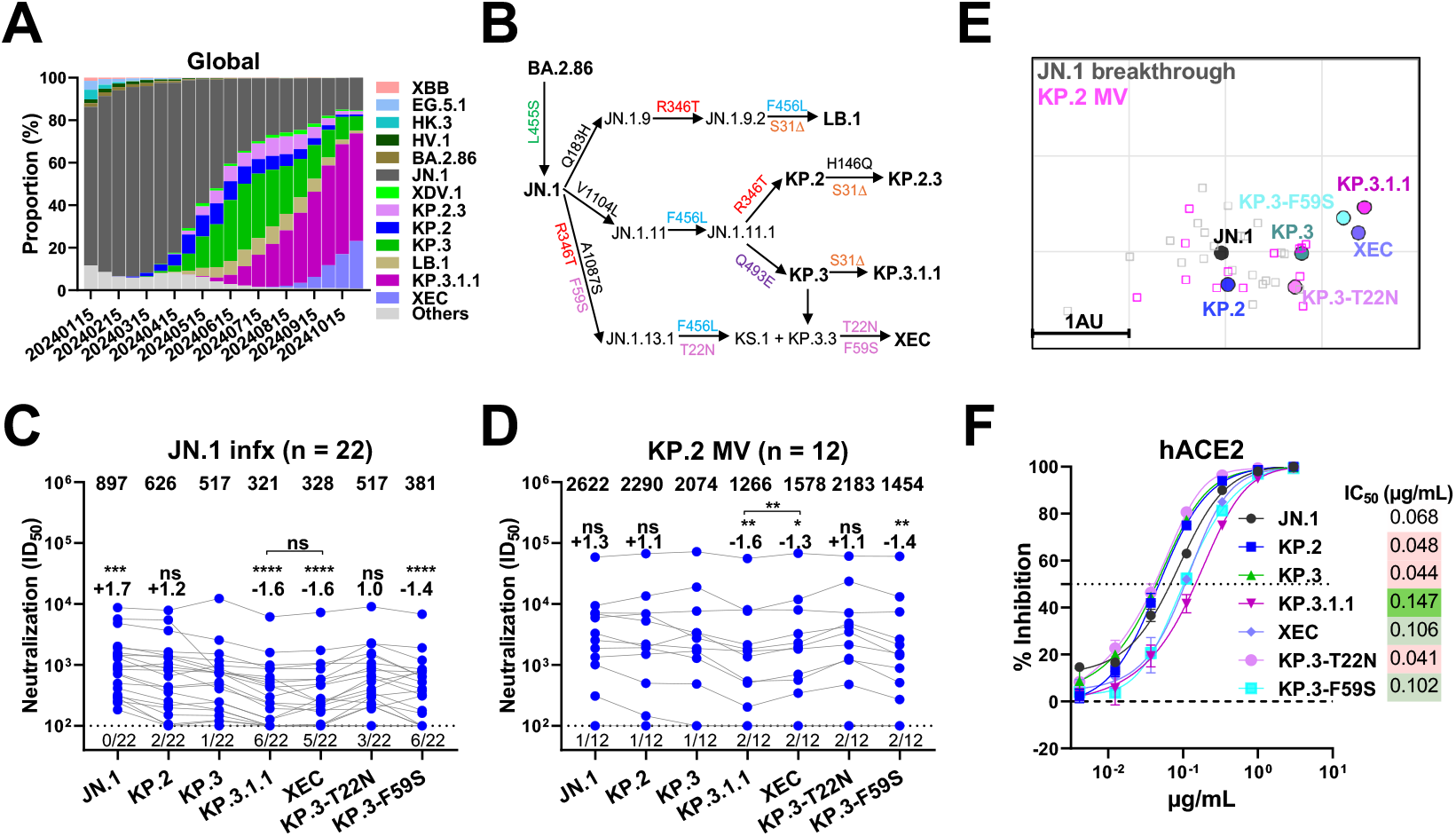
Characterization of SARS-CoV-2 JN.1 sublineages. **A**. Relative frequencies of dominant SARS-CoV-2 subvariants from 2024/01 to 2024/10; data from GISAID^1^. **B**. Viral evolutionary pathways and spike mutations of the indicated JN.1 subvariants. **C-D**. Serum neutralizing titers (ID_50_) against VSV-based pseudoviruses bearing spike proteins from SARS-CoV-2 JN.1 sublineages for samples from cohorts: “JN.1 infx” (**C**) and “KP.2 MV” (**D**). Compared with KP.3, KP.3.1.1 carries an S31Δ spike mutation in the NTD. The geometric mean ID_50_ titer (GMT) is presented at the top of the panel. The fold change in GMT for each virus compared to KP.3 is also shown immediately above the symbols. Statistical analyses used Wilcoxon matched-pairs signed-rank tests. n, sample size. ns, not significant; *p < 0.05; **p < 0.01; ***p < 0.001; ****p < 0.0001. Numbers under the dotted lines denote numbers of serum samples that were under the limit of detection (ID_50_ <100). **E**. Antigenic map generated using all neutralization data from panels **C** and **D**. One antigenic unit (AU) represents an approximately 2-fold change in ID_50_ titer. Serum samples and viruses are shown as squares and dots, respectively. **F**. Sensitivity of JN.1 subvariants to hACE2 inhibition. IC_50_ values are also presented. Data are shown as mean ± standard error of mean (SEM) for three technical replicates.

To address this question, we first tested serum neutralization against VSV pseudotyped KP.3.1.1, XEC, and XEC individual spike mutations on KP.3 (KP.3-T22N and KP.3-F59S), compared to JN.1, KP.2, and KP.3, using samples from two cohorts of adults: 1) participants with a history of JN.1 sublineage infection during 2024, sampled 32-87 days after infection (“JN.1 infx”), and 2) participants who received an updated KP.2-based mRNA monovalent vaccine booster, sampled approximately 4 weeks after dosing (“KP.2 MV”) (**Appendix Table S1 and S2**). Compared with KP.3.1.1, XEC demonstrated a similar level of evasion to serum neutralization in the JN.1 infx cohort (**Figure 1C**) but was slightly more sensitive to serum neutralization in the KP.2 MV cohort (**Figure 1D**). KP.3 was 1.3-to-1.7-fold more resistant to serum neutralization than JN.1, while KP.3.1.1 and XEC were 1.3-to-1.6-fold more resistant than KP.3. These increases in resistance to serum neutralization were explained by the component mutations S31Δ and F59S, individually tested on the background of KP.3 as KP.3.1.1 and KP.3-F59S, respectively (**Figure 1C and 1D**). In addition, serum neutralizing titers in KP.2 MV participants were generally higher than in JN.1 infx participants, with levels correlated with clinical protection^2^.

The serum neutralization data from both cohorts were used to generate antigenic maps for these subvariants. We observed that KP.3.1.1, XEC, and KP.3-F59S clustered closely together, approximately 1.2 antigenic units from JN.1, while KP.3 and KP.3-T22N exhibited a shorter but similar antigenic distance to JN.1 (**Figure 1E**). These results indicate that F59S and S31Δ are comparable antigenically, while T22N has minimal impact on neutralization by these sera.

We then performed neutralization assays using a panel of monoclonal antibodies that retained potency against KP.3 and directed to multiple epitopes on the viral spike. S31Δ and F59S knocked out the NTD-SD2-specific antibody C1717^3^ and impaired receptor-binding domain (RBD) class 4/1 antibodies VYD222 (Pemivibart)^4^ and 25F9^5^, potentially explaining the increased resistance of KP.3.1.1 and XEC to serum neutralization. (**Appendix Figure S1A**). But how do mutations at the bottom of NTD affect monoclonal antibodies directed to the top of the spike? To address this question, we tested the inhibition of soluble human ACE2 (hACE2) against the same panel of pseudoviruses. T22N did not alter the susceptibility of KP.3 to hACE2 inhibition, while S31Δ and F59S impaired hACE2 inhibition by 3.3- and 2.3-fold (**Figure 1F**), respectively, indicating a lower affinity for the viral receptor. The above findings collectively suggested that the upward motion of the RBD may be impaired by either S31Δ or F59S, since the spike binding to both the receptor and class 4/1 antibodies requires the RBD to be in the up position. Indeed, structural analysis showed that S31 and F59 interact via hydrogen bonding (**Appendix Figure S1B**), and that S31Δ and F59S mutations were functionally equivalent in altering NTD conformation, not only enabling escape from NTD-SD2-directed antibodies like C1717 but also likely indirectly hindering the upward movement of RBD.

In summary, KP.3.1.1 and XEC demonstrate greater antibody evasion than JN.1 and KP.3, which likely contributes to their increasing global prevalence. The functional equivalence of S31Δ and F59S mutations renders KP.3.1.1 and XEC antigenically similar.

