## Supplementary Appendix for "Antibody evasiveness of SARS-CoV-2 subvariants KP.3.1.1 and XEC"

### Materials and Methods

#### Clinical cohorts

Serum samples were collected as part of the VIVA study at the University of Michigan[1,2] and as part of the "COVID-19 Persistence and Immunology Cohort (C-PIC)" study at Columbia University. Specimens were obtained following participant consent and in adherence to the protocols approved by the Institutional Review Board of the University of Michigan Medical School (protocol HUM00232359) and Columbia University (protocol AAAS9722), respectively.

In this study, serum samples were collected from two cohorts: 1) individuals with a recent JN.1 sublineage infection ("JN.1 infx"); and 2) individuals who had been administered the updated KP.2 monovalent booster ("KP.2 MV"). The majority of the study subjects were female, representing 88.2%, with an average age of 55.3 years. Serum samples were collected, on average, 54.8 days post-JN.1 sublineage infection and 30.5 days post-KP.2 booster. Demographic details, vaccination status, and serum collection timelines are summarized for each cohort in **Table S1 and Table S2**.

#### Cell lines

HEK293T (ATCC, CRL-3216) cells and Vero-E6 cells (ATCC, CRL-1586) were cultured in Dulbecco's modified Eagle's medium (DMEM) supplemented with 10% heat-inactivated fetal bovine serum and 1% penicillin-streptomycin. All cell lines were cultured in an atmosphere of 5% $CO_2$ at 37°C.

#### Construction of SARS-CoV-2 spike plasmids

The spike constructs of JN.1, KP.2, and KP.3 were generated as previously reported[1]. The spike gene of KP.3.1.1 and XEC, as well as spike gene constructs bearing mutations T22N and F59S, were generated using Q5 site-directed mutagenesis with the KLD master mix kit (NEB). All constructs were confirmed by Sanger sequencing.

#### Pseudovirus production

Pseudotyped SARS-CoV-2 was produced following a previously established protocol[3] . HEK293T cells were first transfected with spike-encoding plasmids using 1 mg/mL of PEI-MAX (Polysciences, Inc.) and cultured for 24 hours. The transfected HEK293T cells were then infected with VSV-G pseudotyped ΔG-luciferase virus (Kerafast, EH1020-PM) at a multiplicity of infection (MOI) of approximately 3 to 5. Two hours later, the cells were washed three times with complete culture medium and cultured in fresh medium for another 24 hours. The transfection supernatant was then harvested and clarified by centrifugation at 2000 rpm for 10 minutes. Each viral stock was subsequently incubated with 20% I1 hybridoma (ATCC, CRL-2700) supernatant for 1 hour at room temperature to neutralize contaminating VSV-G particle before measuring titers and making aliquots for storage at -80°C until use.

**Serum and monoclonal antibody neutralization, and ACE2 inhibition assays**

Pseudoviruses were titrated to standardize the viral input prior to each neutralization or inhibition assay. For neutralization assays, serum samples were inactivated at 56°C for 30 minutes before use, and the inactivated sera were diluted by a factor of 100 followed by a series of 7 four-fold serial dilutions. For monoclonal antibody neutralization assays, each antibody was diluted from 10 µg/mL with a dilution factor of five across 7 serial dilutions. For ACE2 inhibition assays, as reported in our study[4], soluble chimeric human ACE2 (hACE2), which contains ACE2 residues 1-732 and is fused to human IgG1 Fc, was diluted from 3 µg/mL with a dilution factor of three across 7 serial dilutions. Following this, pseudoviruses were added and incubated at 37 °C for 1 hour. As a control, wells containing only the pseudovirus were also prepared on each test plate. Subsequently, Vero-E6 cells were seeded at 40,000 cells per well and were incubated overnight at 37°C on either neutralization or inhibition test plates. Afterwards, cellular lysis was conducted, and the resultant luciferase activity was quantified employing the Luciferase Assay System (Promega) in tandem with Tecan Infinite® 200 PRO using i-control™ software v.3.9.1.0, in accordance with the manufacturer's instructions. The serum dilution that inhibits 50% of virus entry ($ID_{50}$), or the half-maximal inhibitory concentration ($IC_{50}$) by antibody and hACE2, was calculated using nonlinear five-parameter dose-response curve fitting using GraphPad Prism v.10.3.

**Antigenic cartography**

Antigenic cartography for the JN.1 subvariants was conducted through the integration of $ID_{50}$ titers from individual sera, as previously described[4-6]. The visual representations were generated utilizing the Racmacs package (version 1.1.4, accessible at https://acorg.github.io/Racmacs/) within the R computational environment, version 4.0.3. The algorithmic optimization process was executed over 2,000 iterations, with the 'minimum column basis' set to 'none'.

**Quantification and statistical analysis**

$ID_{50}$ values for serum neutralization and $IC_{50}$ values for monoclonal antibody neutralization and hACE2 inhibition were obtained from a five-parameter dose-response curve using GraphPad Prism v.10.3. Statistical analyses were conducted by Wilcoxon matched-pairs signed-rank tests in the same software. Levels of statistical significance are annotated as: ns, not significant; $*p < 0.05$; $**p < 0.01$; $***p < 0.001$ and $****p < 0.0001$.

**Acknowledgements**

This study was supported by funding from the NIH SARS-CoV-2 Assessment of Viral Evolution (SAVE) Program (subcontract no. 0258-A700-4609 under federal contract no. 75N93021C00014 to D.D.H. and (subcontract GR0010139-PO024016 under federal contract no. 75N93021C00016) to A.G. and the Gates Foundation (project INV019355) to D.D.H., internal startup funding UR014016 from Columbia University to Y.G. K23 AI171263 to L.J.P., K24 AI155230 to M.T.Y. We thank all who contributed their data to the Global Initiative on Sharing All Influenza Data (GISAID).

We express our gratitude to Jayesh Shah, Amanda Castillo, Meredith McNairy and Antonia Sturizo for conducting the C-PIC study (Columbia), and to Zijin Chu, Theresa Kowalski-Dobson, Anna Buswinka, Gabe Simjanovski, Joseph Wendzinski, Mayurika Patel, Kathleen Lindsey, and Dawson Davis of the VIVA study team for conducting the VIVA study.

**Author contributions**

The study was conceptualized by Q.W., Y.G., I.A.M., and D.D.H. Experiments were conducted and data analyzed by Q.W., Y.G., I.A.M., and M.W. Project management was handled by Q.W. Serum samples were collected by H.M., C.G., R.V., L.J.P., M.T.Y., A.G., and their colleagues. The results were analyzed, and the manuscript was written by Q.W., Y.G., M.W., I.A.M., and D.D.H. All contributing authors have reviewed and endorsed the manuscript.

**Declaration of interests**

D.D.H. co-founded TaiMed Biologics and RenBio, and he serves as a consultant for WuXi Biologics and Brii Biosciences and is a board director at Vicarious Surgical. A.G. served as a member of the scientific advisory board for Janssen Pharmaceuticals and has consulted and serves on a scientific advisory board for Sanofi Pasteur. The remaining authors declare no conflicts of interest.

**Table S1: Summary of clinical cohorts.**

infx, infection; MV, monovalent vaccine; WT, wildtype; BV, bivalent vaccine.

| | | All participants | | JN.1 infx | | KP.2 MV | |
|---|---|---|---|---|---|---|---|
| | | No. or Mean | % or (range) | No. or Mean | % or (range) | No. or Mean | % or (range) |
| **Total** | | 34 | | 22 | | 12 | |
| **Female** | | 30 | 88.2% | 19 | 86.4% | 11 | 91.7% |
| **Male** | | 4 | 11.8% | 3 | 13.6% | 1 | 8.3% |
| **Age** | | 55.3 | (24, 81) | 55.9 | (33, 78) | 54.5 | (24, 81) |
| **No. Vaccines** | All vaccines | 5.7 | (2, 9) | 5.1 | (2, 9) | 6.8 | (6, 9) |
| | WT | 3.5 | (2, 5) | 3.4 | (2, 4) | 3.8 | (3, 5) |
| | BA.5 BV | 0.9 | (0, 2) | 0.9 | (0, 2) | 1.0 | (0, 1) |
| | XBB.1.5 | 0.9 | (0, 2) | 0.85 | (0, 2) | 1.0 | (0, 2) |
| | KP. 2 MV | 0.4 | (0,1) | 0 | (0,0) | 1 | (1,1) |
| **No. Infections** | | 1.4 | (0, 3) | 1.6 | (1, 3) | 0.8 | (0, 2) |
| **Sera Days Post Infection** | | 155.4 | (0, 835) | 54.8 | (32, 87) | 598.4 | (0, 835) |
| **Sera Days Post Vaccination** | | 230.0 | (20, 1346) | 350.6 | (20, 1346) | 30.5 | (25, 36) |

**Table S2: Participant demographic, vaccine, and infection details.** Vaccine formulations are denoted as wildtype (WT), BA.5 Bivalent (BA.5), XBB.1.5 monovalent (XBB.1.5), and KP.2 monovalent (KP.2). Vaccine manufacturers are denoted as Pfizer (P), Moderna (M), Janssen (J), Other (O), and Unknown (U). Yr, years; Infx, infection; Vax, vaccination; DPI, days post infection; DPV, days post vaccination.

| ID | Age (Yr) | Sex | Race | No. Infx | No. Vax | DPI | DPV | Vaccine History |
|---|---|---|---|---|---|---|---|---|
| **JN.1 infx** | | | | | | | | |
| JN.1-11 | 71 | M | Asian | 2 | 5 | 40 | 315 | WT-P/WT-P/WT-J/BA.5-P/XBB.1.5-P |
| JN.1-12 | 52 | F | Asian | 2 | 3 | 39 | 327 | WT-P/WT-P/XBB.1.5-P |
| JN.1-13 | 66 | F | White | 2 | 6 | 59 | 151 | WT-P/WT-P/WT-P/WT-P/BA.5-P/XBB.1.5-M |
| JN.1-14 | 35 | F | White | 2 | 5 | 32 | 155 | WT-P/WT-P/WT-P/BA.5-P/XBB.1.5-P |
| JN.1-15 | 49 | F | White | 2 | 5 | 35 | 127 | WT-P/WT-P/WT-M/BA.5-P/XBB.1.5-P |
| JN.1-16 | 38 | F | White | 2 | 5 | 39 | 162 | WT-P/WT-P/WT-P/BA.5-P/XBB.1.5-M |
| JN.1-17 | 78 | F | White | 3 | 6 | 77 | 169 | WT-P/WT-P/WT-P/WT-P/BA.5-P/XBB.1.5-P |
| JN.1-18 | 33 | F | White | 1 | 4 | 81 | 203 | WT-P/WT-P/WT-P/XBB.1.5-P |
| JN.1-19 | 71 | F | White | 1 | 9 | 80 | 20 | WT-M/WT-M/WT-M/WT-M/WT-M/BA.5-M |
| JN.1-20 | 67 | F | Asian | 2 | 7 | 54 | 25 | WT-P/WT-P/WT-P/WT-P/BA.5-P/XBB.1.5-M |
| JN.1-21 | 62 | F | White | 2 | 6 | 55 | 117 | WT-P/WT-P/WT-P/WT-M/BA.5-M/XBB.1.5-P |
| JN.1-22 | 48 | F | White | 1 | 3 | 55 | 847 | WT-P/WT-P/WT-P |
| JN.1-23 | 69 | F | White | 2 | 3 | 55 | 847 | WT-P/WT-P/WT-P |
| JN.1-24 | 41 | F | Asian | 2 | 5 | 87 | 179 | WT-P/WT-P/WT-P/BA.5-M/XBB.1.5-M |
| JN.1-25 | 65 | F | White | 1 | 4 | 65 | 218 | WT-J/WT-J/BA.5-P/XBB.1.5-P |
| JN.1-26 | 50 | M | White | 2 | 2 | 67 | 1346 | WT-P/WT-P |
| JN.1-27 | 78 | M | White | 1 | 7 | 39 | 191 | WT-P/WT-P/WT-P/WT-P/BA.5-P/BA.5-P |
| JN.1-28 | 57 | F | White | 1 | 5 | 61 | 608 | WT-P/WT-P/WT-P/BA.5-M/BA.5-P |
| JN.1-29 | 59 | F | White | 2 | 6 | 36 | 188 | WT-M/WT-M/WT-M/WT-M/BA.5-P/XBB.1.5-M |
| JN.1-30 | 64 | F | White | 1 | 6 | 40 | 253 | WT-P/WT-P/WT-P/WT-P/BA.5-P/XBB.1.5-P |
| JN.1-31 | 42 | F | White | 1 | 5 | 70 | 192 | WT-P/WT-P/WT-P/BA.5-M/XBB.1.5-O |
| JN.1-32 | 34 | F | Other | 1 | 3 | 39 | 1013 | WT-P/WT-P/WT-P |
| **KP.2 MV** | | | | | | | | |
| KP.2-V1 | 24 | F | White | 1 | 6 | 835 | 28 | WT-P/WT-P/WT-P/BA.5-M/XBB.1.5-P/KP.2-P |
| KP.2-V2 | 25 | F | White | 0 | 6 | - | 32 | WT-P/WT-P/WT-M/BA.5-M/XBB.1.5-M/KP.2-P |
| KP.2-V3 | 81 | F | White | 1 | 7 | 662 | 31 | WT-P/WT-P/WT-P/WT-M/BA.5-P/XBB.1.5-M/KP.2-M |
| KP.2-V4 | 74 | F | White | 2 | 7 | - | 33 | WT-P/WT-P/WT-P/WT-P/BA.5-P/XBB.1.5-M/KP.2-P |
| KP.2-V6 | 33 | F | White | 2 | 6 | 335 | 35 | WT-P/WT-U/WT-P/BA.5-P/XBB.1.5-P/KP.2-P |
| KP.2-V7 | 75 | F | Asian | 0 | 6 | - | 36 | WT-M/WT-M/WT-M/WT-M/WT-M/KP.2-M |
| KP.2-V8 | 59 | F | White | 0 | 7 | - | 35 | WT-P/WT-P/WT-P/WT-P/BA.5-P/XBB.1.5-M/KP.2-P |
| KP.2-V9 | 68 | F | White | 0 | 9 | - | 33 | WT-P/WT-P/WT-P/WT-P/WT-P/BA.5-P/XBB.1.5-P/XBB.1.5-P/KP.2-P |
| KP.2-V10 | 78 | F | White | 0 | 9 | - | 31 | WT-P/WT-P/WT-P/WT-M/BA.5-M/BA.5-M/XBB.1.5-M/XBB.1.5-M/KP.2-M |
| KP.2-V11 | 35 | M | Asian | 1 | 4 | - | 25 | WT-P/WT-P/WT-P/KP.2-M |
| KP.2-V12 | 25 | F | Asian | 0 | 6 | - | 25 | WT-P/WT-P/WT-P/BA.5-M/XBB.1.5-M/KP.2-M |
| KP.2-V13 | 70 | F | Asian | 3 | 8 | 585 | 17 | WT-P/WT-P/WT-P/WT-P/BA.5-P/BA.5-P |

**A**

| $IC_{50}$ (μg/mL) | | JN.1 | KP.2 | KP.3 | KP.3.1.1 | XEC | KP.3-T22N | KP.3-F59S |
|---|---|---|---|---|---|---|---|---|
| **NTD-SD2** | **C1717** | 1.485 | 0.669 | 1.084 | >10 | >10 | 1.563 | >10 |
| **RBD class 1** | **BD55-1205** | 0.005 | 0.002 | 0.001 | 0.003 | 0.002 | 0.001 | 0.002 |
| | **BD55-4637** | 0.024 | 0.022 | 0.016 | 0.045 | 0.032 | 0.016 | 0.035 |
| **RBD class 3** | **CYFN1006-1** | 0.004 | 0.009 | 0.003 | 0.003 | 0.003 | 0.004 | 0.004 |
| **RBD class 4/1** | **SA55** | 0.003 | 0.003 | 0.002 | 0.003 | 0.003 | 0.002 | 0.003 |
| | **VYD222** | 0.096 | 0.121 | 0.415 | 2.675 | 1.753 | 0.512 | 2.009 |
| | **25F9** | 0.780 | 0.862 | 0.966 | 4.366 | 2.500 | 0.589 | 2.183 |
| **RBD-NTD interface** | **C68.61** | 0.677 | 0.456 | 0.537 | 0.998 | 1.025 | 0.320 | 0.776 |

**B**

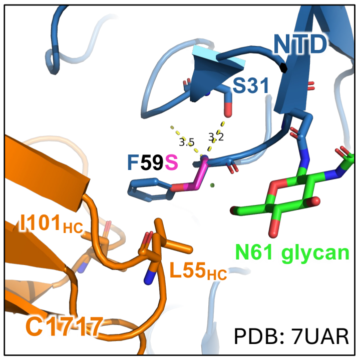

**Figure S1: Monoclonal antibody neutralization of the indicated JN.1 sublineage variants and structural analysis.**

**A.** Monoclonal antibody neutralization against the indicated pseudoviruses. Antibody concentrations resulting in 50% inhibition of infectivity ($IC_{50}$) are presented.

**B.** Structural analysis of S31Δ and F59S in the complex of C1717 and NTD.
